# Development of a Novel Air-Liquid Interface Airway Tissue Equivalent Model for In Vitro Respiratory Modeling Studies

**DOI:** 10.1101/2022.09.21.508886

**Authors:** Timothy Leach, Uma Gandhi, Kimberly D. Reeves, Kristina Stumpf, Kenichi Okuda, Frank C. Marini, Steve Walker, Jeannie Chan, Laura A. Cox, Anthony Atala, Sean V. Murphy

## Abstract

The human airways are complex structures with important interactions between cells, extracellular matrix (ECM) proteins and the biomechanical microenvironment. A robust, well-differentiated *in vitro* culture system that accurately models these interactions would provide a useful tool for studying normal and pathological airway biology. Here, we report the feasibility and analysis of a physiologically relevant air-liquid interface (ALI) 3D airway ‘organ tissue equivalent’ (OTE) model with three novel features: native pulmonary fibroblasts, solubilized lung ECM, and hydrogel substrate with tunable stiffness and porosity. We demonstrate the versatility of the OTE model by evaluating the impact of these features on human bronchial epithelial (HBE) cell phenotype. Variations of this model were analyzed during 28 days of ALI culture by evaluating epithelial confluence, trans-epithelial resistance, and epithelial phenotype via multispectral immuno-histochemistry and next-generation sequencing. Cultures that included both solubilized lung ECM and native pulmonary fibroblasts within the hydrogel substrate formed well-differentiated ALI cultures that maintained a barrier function and expressed mature epithelial markers relating to goblet, club and ciliated cells. Modulation of hydrogel stiffness did not negatively impact HBE differentiation and could be a valuable variable to alter epithelial phenotype. This study highlights the feasibility and versatility of a 3D airway OTE model to model the multiple components of the human airway 3D microenvironment.

## Introduction

The human bronchial tree is a complicated heterogeneous system that has important functions beyond being a simple barrier and conduit for air exchange. The airways are at the interface of the internal and external environment of the human body facilitating a variety of functions including mucociliary clearance, airway humidification, pathogen/particulate sensing and defense, and signaling to the underlying mesenchyme and immune system^1,2^. The airway epithelium is heterogeneous with distinct specialized cells including ciliated cells, goblet cells, club cells, basal cells, ionocytes, and neuroendocrine cells^2^. For many diseases there is dysregulation in these epithelial cell subtypes that directly corresponds with the disease including asthma, chronic obstructive pulmonary disorder, and cystic fibrosis^3,4^. Additionally, there is significant interplay with the pseudostratified epithelium and its basal three-dimensional (3D) microenvironment that includes subepithelial fibroblasts, immune cells, endothelial cells, and smooth muscle cells depending on airway size. Each of these cells found in the interstitial layer of the airways play key roles in the cell-cell communication that influence normal function and disease^5-7^. Besides the heterogenic cell populations within the airways, the subepithelial extracellular matrix (ECM) has shown to play a key role in airway structural integrity as well as in cell regulatory functions such as cell activation, proliferation and differentiation^8^. These proteins and polymers are tissue-specific and provide specific ECM-cell receptor signaling. For instance, the composition of collagen within the airways along with the corresponding stiffness has been directly correlated to diseased phenotypes and pathologically activated cells^9^. Despite advances in pulmonary medicine, we currently lack an optimal non-clinical model comprised of these important cell-cell, ECM-cell, and biomechanical components of the human airways.

The current gold standard for the study of the human airways is a monoculture of human bronchial epithelial (HBE) cells cultured on porous polymer membranes at an air-liquid interface (ALI) ^1,10^. The main advantage of this model is the air interface which promotes HBE differentiation and pseudostratification. The ALI allows for mucus production and ciliation of the HBE as well as many other important functions such as barrier properties, aerosolized exposure, wound healing and regeneration. In conjunction with *in vivo* animal models, this model has provided many years of successful application to better understand normal airway epithelial function, disease, injury and evaluation of therapeutics^11-13^. Still, this 2D ALI model has limitations, including: 1) lack of physiological cell-cell interactions with non-epithelial cells, such as the underlying stromal cells, 2) lack of cell-ECM matrix interactions beyond collagen coating of the polymer surface to mimic the basal membrane, and 3) a growth surface that is orders of magnitude stiffer than the native *in vivo* tissue microenvironment. These non-physiological conditions are likely to impact HBE phenotype, heterogeneity, and functionality *in vitro* resulting in an imprecise representation of human airway epithelial phenotype, altered pharmacodynamics, and limitations with modeling complex disease^1,14-20^. The absence of a 3D environment affects its potential for studying diseased bronchial conditions due to the complex interactions of most conditions that involve the surrounding tissue environment^21,22^.

Therefore, there has been significant interest in the development of 3D cell culture models and microfluidic models to address these limitations and better represent the microenvironment experienced by cells *in vivo*. There have been two main approaches to date: spheroidal models and 2D planar microfluidic models. Spheroidal models address some limitations of 2D monoculture models by providing improved cell-cell interactions along with the possibility of an ECM-based microenvironment. A major drawback of the spheroidal design though is the lack of an external air liquid interface for analysis of the cilia and mucus along with aerosolized drug and toxin exposure. 2D planar microfluidic models have been developed to allow for physiological air and liquid flow on the apical and basal sides of the HBEs, respectively^23-25^. Some of these 2D planar microfluidic models include co-cultures of HBEs with other cell types, often plated on the underside of the polymer membrane or in parallel membranes within the same microfluidic chip ^23,25,26^. A major limitation of this design is the lack of a biomechanical ECM environment beyond collagen coating of the stiff polymer membrane surface to support cell attachment. Therefore, there remains a critical need in the field for a 3D *in vitro* airway model that allows for both the formation of well-differentiated HBE cultures at ALI and a 3D microenvironment that includes physiological cell-cell, ECM-cell and biomechanical interactions.

Here we report the development of a planar airway 3D organ tissue equivalent (OTE) model, comprised of a well-differentiated HBE layer at ALI, maintained on a hydrogel substrate layer that can contain native lung fibroblasts and lung sECM (**Fig. 1**). The interstitial hydrogel layer is composed of thiolated HA, thiolated gelatin, and a polyethylene glycol (PEG) crosslinker that are photocrosslinkable for spatiotemporal control of the biogel’s biomechanical properties. This system can be fabricated on standard transwells of any size or membrane pore size, or incorporated into microfluidic platforms^27,28^. We evaluated the incorporation of decellularized and solubilized human lung ECM (sECM) and native pulmonary fibroblasts into the hydrogel layer, and compared a range of hydrogel stiffnesses representing physiological values expected in airway tissue. By varying the PEG crosslinker composition we can alter the elastic modulus of the hydrogels within a physiologically representative range of airway stiffnesses for healthy and pathological bronchial tissue without affecting the ECM composition of the hydrogel ^29,30^. Meanwhile, the incorporation of human lung sECM and native lung fibroblasts are likely to provide airway-specific cytokine and ECM signaling molecules that improve HBE differentiation and attachment^31,32^. These model variants were characterized by measuring HBE growth, functionality, and differentiation compared to 2D ALI cultures and native human airway tissue samples.

**Figure 1.**
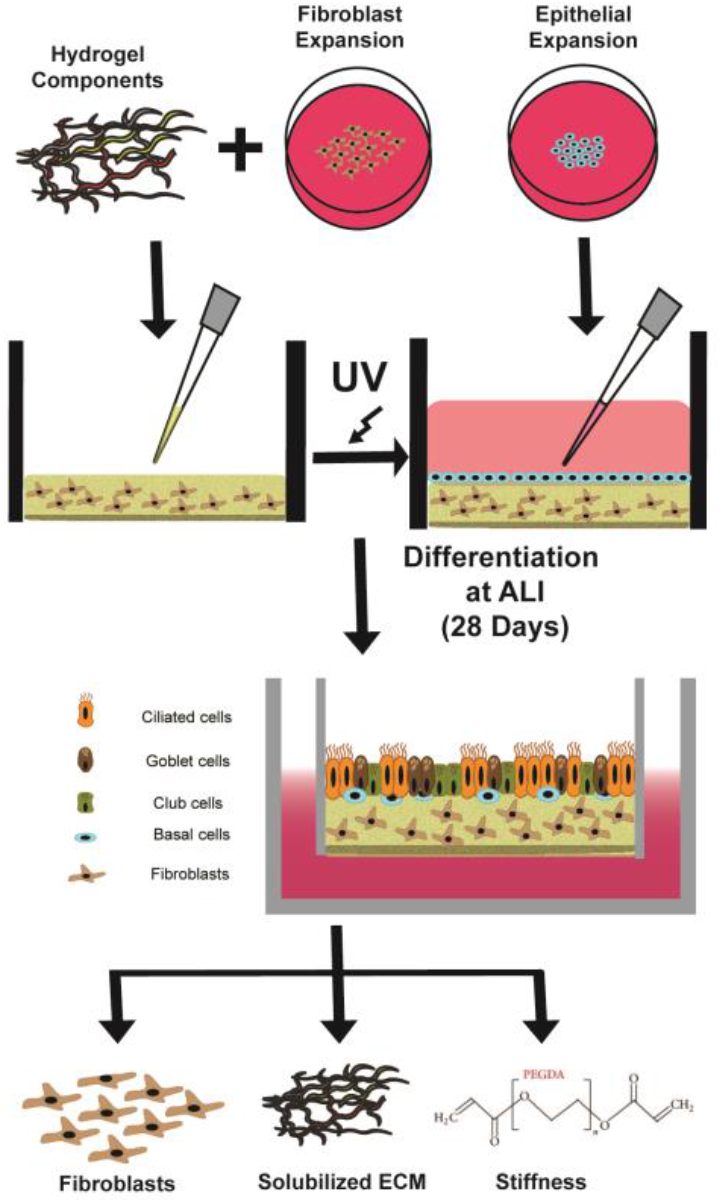
Organ Tissue Equivalent Fabrication Schematic. To fabricate the complete 3D OTE model, native lung fibroblasts were mixed with the hydrogel components and solubilized lung ECM and 80μL was pipetted into an 8μm polycarbonate membrane Transwell. After UV crosslinking, the HBEs were cultured on the apical side and allowed to differentiate at ALI for 28 days. The three main components of interest for the final OTE model are the native lung fibroblasts, solubilized lung ECM, and biomimetic hydrogel with stiffness adjusted via different crosslinkers.

## Results

### SOLUBILIZED LUNG EXTRACELLULAR MATRIX CHARACTERIZATION

Organ ECM composition has shown to be tissue-specific^31^, and the composition could provide insight into crucial ECM components for well-differentiated HBEs that interact via cell adhesion and cell-ECM receptor signaling. We integrated decellularized lung sECM into the hydrogel component of our 3D OTE model, and evaluated its impact on HBE viability and a differentiated phenotype. A total of six healthy human lungs were decellularized and processed for analysis to provide a general characterization of the lung ECM microarchitecture (**Fig. 2A**). Complete decellularization was confirmed with no discernible nuclei or other cellular components after H&E staining (**Suppl. Fig. S1**). The sECM solution was quantified with assays for total protein and a variety of major ECM components to evaluate retention of these important components. After solubilization, between 3.3-5.4mg/mL of total protein was recovered (**Fig. 2B**). We also quantified individual ECM components: collagen (338.8 ± 116.5μg/mL), elastin (241.7 ± 111.9μg/mL), laminin (9.80 ± 2.48μg/mL), fibronectin (70.4 ± 21.1μg/mL), sulfated glycosaminoglycans (sGAGs) (195.5 ± 103.4μg/mL) and the major non-sulfated GAG, hyaluronic acid (8.2 ± 4.3μg/mL) (**Fig. 2C**). Of the total collagen recovered, the majority of the collagen analyzed, based on spectral counts, was from the following types: type I (17.2 ± 3.2%), type V (36.4 ± 6.7%), and Type VI (21.7 ± 5.6%). After decellularization and solubilization, our human lung sECM retained multiple important ECM components known to be important for HBE growth, attachment and differentiation^33^. The highest concentration of lung sECM (∼2mg/mL of protein) was utilized for fabricating hydrogels that would allow for dissolution of the individual hydrogel components for all subsequent experiments in order to best mimic the ECM environment of the airways.

**Figure 2.**
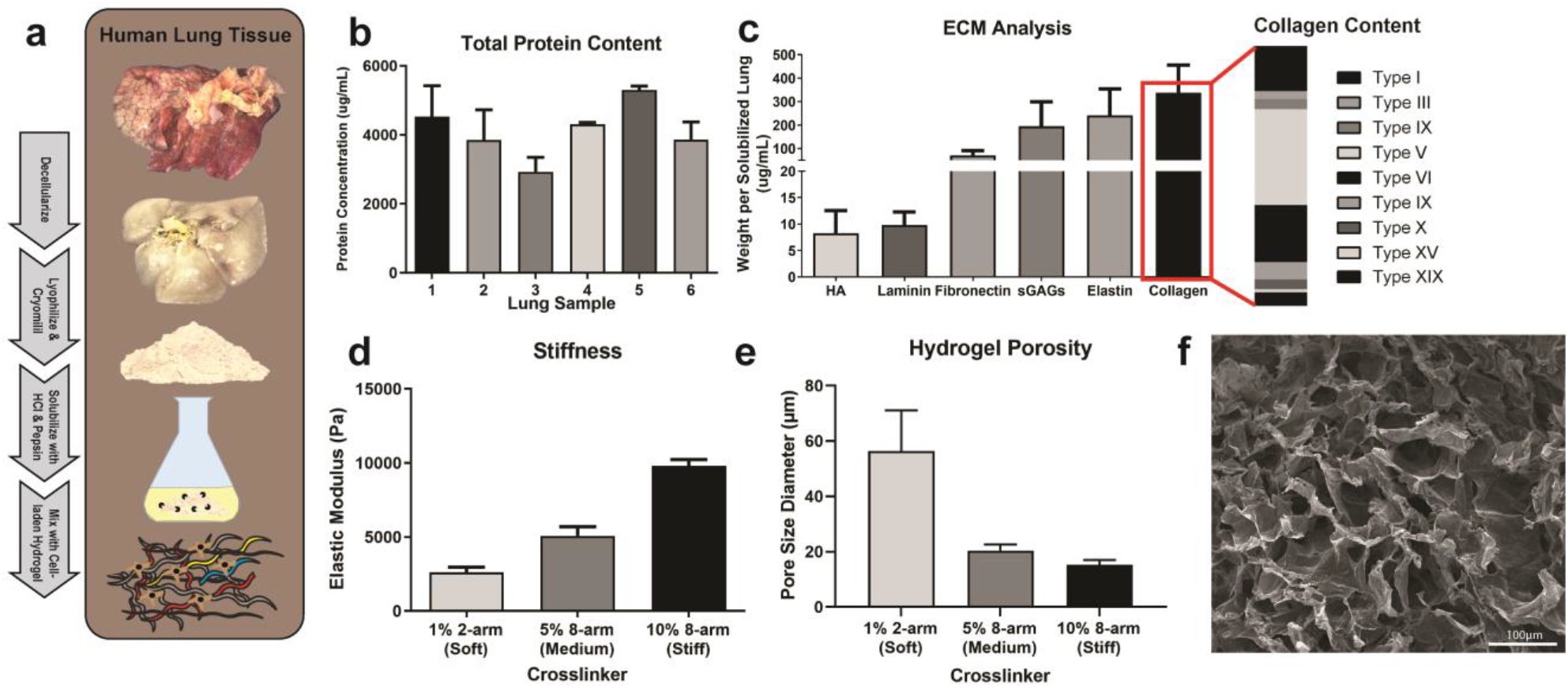
Fabrication & Analysis of Lung Extracellular Matrix-Derived Biogel. **(A)** To obtain the lung ECM biogel, human lung tissue is decellularized, lyophilized and cryomilled, digested, and mixed with a hyaluronic acid and gelatin-based hydrogel to form the biogel. **(B-C)** Utilizing a BCA Protein Assay kit, ECM colorimetric kits, and ECM ELISAs the total amount of protein along with certain ECM components were quantified for each lung sample. The ECM biogel can employ crosslinking molecules of different sizes and geometries to tune the biomechanical properties of the hydrogel, including elastic modulus **(D)** and pore size **(E)**, measured via SEM imaging **(F)**.

### HYDROGEL BIOMECHANICS CHARACTERIZATION

The biomechanical properties of individual tissues have a significant influence on healthy and pathological cell populations and phenotypes. Substrate stiffness directly correlates with attachment of adherent cells and influences cell phenotype^34^. For airway tissue, measurements of airway biomechanical properties report a stiffness range of 1-20kPa depending on bronchi generation and disease status^29,30,35^. For our study, hydrogels were fabricated to model healthy bronchi airway tissue stiffnesses with three hydrogel stiffness groups representing conditions we defined here as: Soft (2.6kPa), Medium (5.1kPa), and Stiff (9.8kPa) (**Fig. 2D**). With an increase in stiffness, there was a corresponding decrease in porosity of the hydrogel as measured by SEM imaging and quantification with average pore sizes of: 56.45±14.65μm (Soft), 20.47±2.19μm (Medium), and 15.20±1.82μm (Stiff) (**Fig. 2E,F**). There was no significant difference on the stiffness of the hydrogel if sECM was included in the hydrogel mixture (**Suppl. Fig. S2**). Increasing elastic modulus also correlated with increased storage modulus between the three stiffness groups (**Suppl. Fig. S3**). The ability to directly influence the hydrogel biomechanical properties without altering ECM concentrations provides a valuable tool for assessing their influence on HBE phenotype when cultured at ALI.

### INFLUENCE OF 3D OTE PARAMETERS ON HBE LAYER

To assess the impact of the individual components incorporated into the OTE model on HBE differentiation and functionality, we fabricated OTEs with hydrogels without either fibroblasts (FB) or lung sECM, with each individually incorporated into the hydrogel, and with both fibroblasts and lung sECM. The OTEs were fabricated with a density of 250,000 fibroblasts/OTE and a lung sECM protein concentration of 2mg/mL to evaluate the feasibility of the OTE model. These conditions were replicated across all three stiffness groups, as summarized below in **Table 1**. HBEs were cultured on the surface of each of these hydrogel groups, airlifted after 4 days, and assessed for differentiation at ALI 28. Each of these OTE groups were evaluated and compared to standard 2D ALI cultures and native human airway for all relevant and compatible analysis. Future studies will expand upon this design by evaluating concentration ranges for FB and lung sECM.

**Table 1.** Experimental Groups Numbered experimental groups for the comparison of 2D ALI culture and the variety of 3D OTE models.

| EXPERIMENTAL GROUPS (n=24) |  |  |  |  |
| --- | --- | --- | --- | --- |
|  | FB+/ECM+ | FB+ | ECM+ | Neither |
| Soft (~2kPa) | **Group 1** | **Group 2** | **Group 3** | **Group 4** |
| Medium (~5kPa) | **Group 5** | **Group 6** | **Group 7** | **Group 8** |
| Stiff (~10kPa) | **Group 9** | **Group 10** | **Group 11** | **Group 12** |
| 2D Culture | **Group 13** |  |  |  |

### BRONCHIAL EPITHELIAL INTEGRITY AND BARRIER FUNCTION

A critical characteristic of ALI cultures is the epithelial barrier that functions to maintain an air-liquid interface and epithelial functionality. The standard assay to assess epithelial integrity and the corresponding polarization is measurement of the trans-epithelial electrical resistance (TEER). Epithelial integrity and barrier function was assessed by TEER and corresponding brightfield whole mount imaging to assess confluency of the epithelial monolayer (**Fig. 3**). We hypothesized that the presence of native FB and sECM would promote HBE attachment and subsequent development of tight junctions for an epithelial barrier.

**Figure 3.**
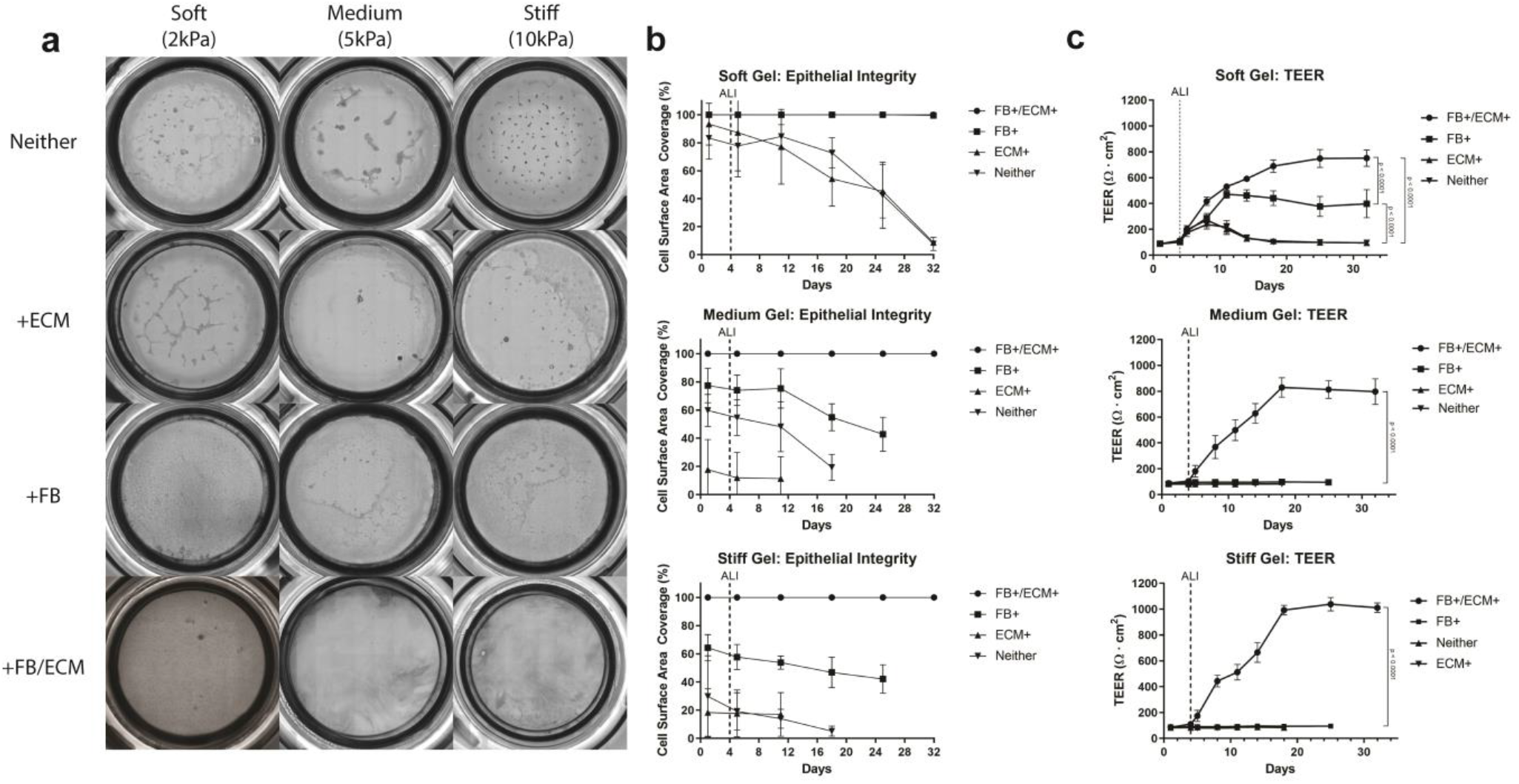
Epithelial Barrier Functionality Analysis. **(A)** Representative whole mount images of the epithelial surface of the OTE cultures during the culture period. **(B)** Epithelial confluency was quantified and plotted for the duration of the culture. **(C)** Trans-epithelial electrical resistance (TEER) of 3D OTE HBE cells during the entire culture period. Significance compared the final day of TEER measurements and the vertical dotted line signifies the day the cultures were changed to ALI. (Note: Some error bars are too small to be shown)

Epithelial detachment and holes in the HBE monolayer could be observed in all OTE cultures that did not include both sECM and lung fibroblasts except for the FB+ only group in the soft hydrogel group (Group 2) (**Fig. 3A**). The image quantification of the epithelium confluency illustrated >99% confluency for these groups compared to all other groups which had <50% confluency (**Fig. 3B**). Qualitatively, the cultures that were unable to maintain a confluent monolayer had HBE cells that either formed aggregates or spread out on the surface in contrast to their typical cobblestone phenotype (**Suppl. Fig. S4)**. We expect that the inclusion of lung sECM along with the influence of native fibroblasts on the ECM microarchitecture, and potentially via production of supportive paracrine factors, provided a more favorable environment for cell attachment and basement membrane generation for a complete HBE monolayer.

TEER measurements supported the image quantification results with the cultures that did not include both FB and sECM dropping to near baseline resistance values at the end of their culture periods (**Fig. 3C**). In contrast, OTEs with both FB and sECM (Groups 1,5,9) had final TEER values more than 7X their baseline values indicating quality epithelial integrity and polarization compared to the other groups (p<0.0001). The soft hydrogel with only FB (Group 2) did show a 4X increase in TEER from baseline but did not develop resistance values as large as the group with both FB and sECM (p<0.0001). An important feature that was noted during the culture period of the OTEs was that none of the hydrogel models contracted from the edges resulting in the loss of the air-liquid interface and barrier function (**Fig. 3A**) typically seen with hydrogels with embedded fibroblasts.

TEER measurements were also taken for the 2D culture and similarly showed a 10X increase from its baseline value at the end of 28 days of ALI culture (**Suppl. Fig. S5**). When compared to the groups that did develop a quality monolayer barrier, the final TEER value of the 2D culture was significantly higher than all of the 3D OTE cultures (p<0.001). This discrepancy is expected to be an effect of measuring electrical resistance through a hydrogel substrate, as imaging showed the widespread presence of ZO-1 tight junction markers in both the 2D and 3D OTE culture (**Suppl. Fig. S6**). Still, there is no *in vivo* standard of electrical resistance measurements for comparison to determine what an ideal physiological electrical resistance is for HBEs.

### HISTOLOGICAL ANALYSIS OF EPITHELIAL PSEUDOSTRATIFICATION AND DIFFERENTIATION

Once OTE hydrogel conditions capable of forming confluent ALI cultures were established representing FB+/ECM+ conditions of all three hydrogel stiffness variants, and FB+ of the soft hydrogel variant (Groups 1, 2, 5 and 9), quantification of their capability to differentiate and HBE phenotype characterization was performed via histological, multispectral immunohistochemical and RNA sequencing transcriptomic analysis. First, we performed H&E staining on the histologically processed cultures collected at the end of ALI culture and compared them to human lower and upper airway tissue samples (**Fig. 4A**). Despite successful cultivation of a complete monolayer of HBE cells, not all groups evaluated demonstrated pseudostratification and qualitative signs of HBE differentiation. Unlike the groups that contained both FBs and sECM, the soft hydrogel with FB-only (Group 2) developed into a 2-3 multi-layered epithelium without clear signs of pseudostratification or ciliation as seen in human airway tissue (data not shown). On the other hand, the groups with both FB and sECM qualitatively showed pseudostratification. This was quantified by MATLAB analysis of the height of the HBE layer for 2D cultures, the OTE groups, and human upper and lower airway tissue (**Fig 4B & Suppl. Fig. S7**). When both FB and sECM were included in the OTE, there was an increase in epithelial height that ranged between lower and upper human airway tissue (Group1: 38.40±1.79μm, Group5: 47.31±10.80μm, Group9: 52.22±11.72μm). All three FB+/sECM+ OTE groups with the different stiffnesses developed an epithelial height significantly larger with regards to 2D ALI culture (2D: 22.78±1.41μm, p<0.0001). Moreover, an increase in stiffness correlated with an increase in pseudostratified height. Specifically, when cultured on the Soft hydrogel (Group 1), the epithelial height was more closely matched with lower airway tissue (38.40±1.79μm vs 34.72±7.79μm, p<0. 01) and when cultured on the Stiff hydrogel (Group 9), the epithelial height was more closely matched with upper airway tissue (52.22±11.72μm vs 61.60±10.70μm, p<0.01). While a larger range of stiffnesses need to be evaluated before such conclusions can be made with confidence, a possible explanation for this matching of epithelial height is the correlation of decreasing tissue stiffness with increasing bronchi generation^35^. This relationship provides an opportunity to develop and compare airway OTE models of different hydrogel stiffnesses for specific bronchial regions. Due to the lack of pseudostratification height and obvious signs of ciliated and goblet cells, only the FB+/sECM+ OTE groups were used for multispectral and transcriptomic analysis of the differentiated HBE phenotype.

**Figure 4.**
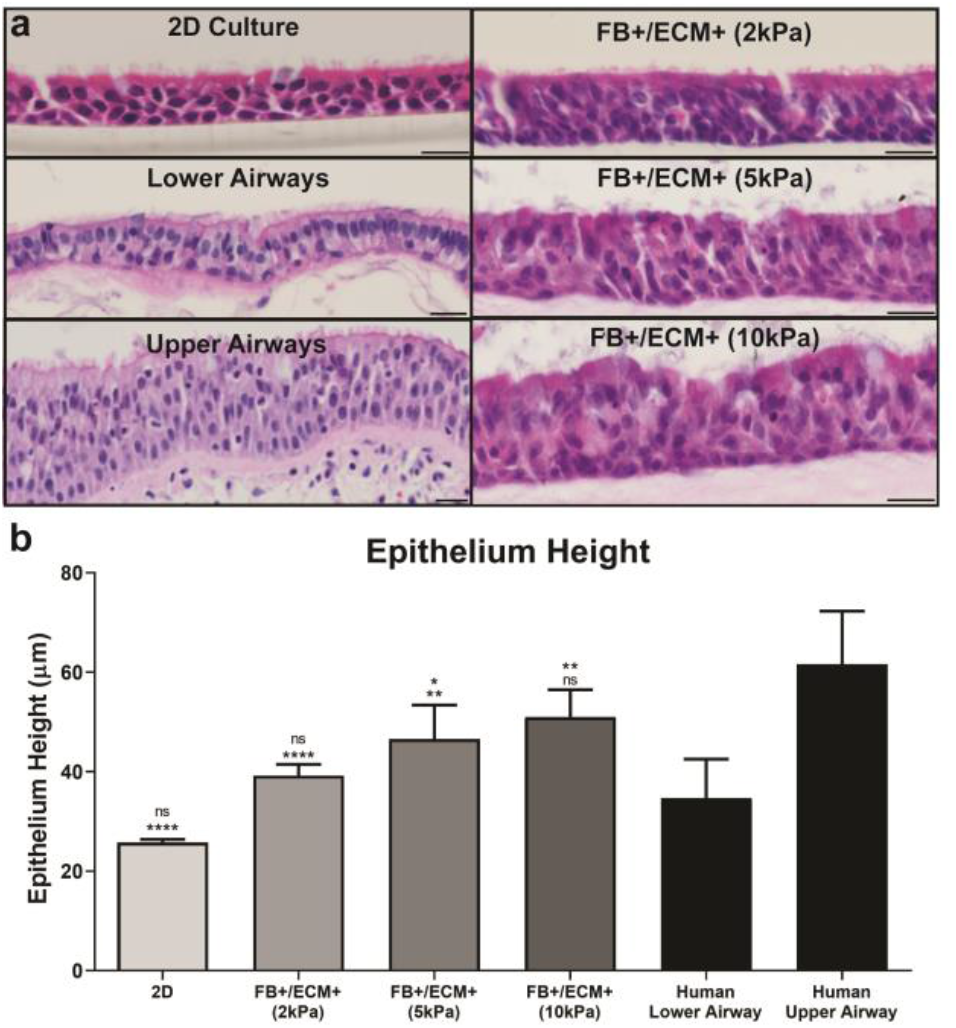
Epithelial Pseudostratification Analysis. **(A)** Representative H&E staining of the FB+/ECM+ OTE model groups, 2D ALI Culture, and upper/lower human airway tissue at ALI day 28. (Scale bar = 20μm). **(B)** Quantitative analysis of the epithelial thickness at ALI day 28 completed via MATLAB image segmentation. The top significance is in relationship to lower airway epithelium and the bottom significance is in relationship to upper airway epithelium (* = p<0.05, ** = p<0.01, **** = p < 0.001).

### MULTISPECTRAL ANALYSIS OF EPITHELIAL DIFFERENTIATION AND FUNCTIONALITY

To characterize the heterogeneous population of differentiated cells in each OTE condition, a Nuance multispectral imaging system was used to quantify multiple molecular markers associated with the different HBE cell subtypes. This system allows for multiplexing and spectral unmixing of images to accurately quantify changes in molecular marker expression between samples while removing autofluorescent noise associated with the tissue, hydrogel, and native ECM. We quantified markers associated with the four major cell types of the airways: MUC5ac (goblet), MUC5b (club), alpha tubulin (ciliated), and TP63 (basal) (**Fig. 5A**). The remaining OTE groups, those that included both sECM and FB at varying stiffness (Groups 1, 5, and 9), and 2D ALI culture were compared against native human airway tissue. When qualitatively compared, all OTE cultures displayed cell type proportions comparable to human airway tissue, while the 2D cultures displayed decreased proportions of each of the evaluated cell types compared to all other groups (**Fig. 5A**). Quantitative analysis of these markers using the Nuance multispectral software demonstrated that all OTE cultures showed a significant increase in staining for each of the evaluated cell types compared to 2D ALI culture (p<0.01) (**Fig. 5B**). OTE cultures had greater MUC5ac staining than 2D cultures (2D: 2.92±0.77, 2kPa: 5.52±1.45, 5kPa: 11.80±2.49, 10kPa: 13.69±2.81), while all groups remained lower compared to human airway tissue (Airways: 25.99±3.68). A similar trend was seen for the club cell marker MUC5b. OTE cultures had greater MUC5b staining than 2D ALI cultures (2D: 17.57±6.58, 2kPa: 47.08±11.59, 5kPa: 63.41±13.30, 10kPa: 63.55±7.37, Airways: 84.04±11.23). Ciliation of the HBE layer showed no statistical difference in ciliated epithelial coverage between human airway tissue and the OTEs, while 2D ALI samples had significantly lower ciliation than airway tissue and OTE groups (2D: 71.11±8.30%, 2kPa: 92.34±6.83%, 5kPa: 91.24±4.65%, 10kPa: 92.81±5.20%, Airways: 85.39±10.24%). Finally, the Medium and Stiff hydrogel groups were statistically similar in TP63 expression compared to airway tissue, while the Soft hydrogel group showed decreased expression, statistically similar to 2D ALI cultures (2D: 12.43±1.31%, 2kPa: 15.15±2.85%, 5kPa: 23.15±6.10%, 10kPa: 25.07±3.77%, Airways: 27.82±3.38%). The quantified increase in each of the evaluated cell-type markers correlated with increasing OTE hydrogel stiffness. These results are indicative of the HBE layer on the OTE models representing a well-differentiated epithelium.

**Figure 5.**
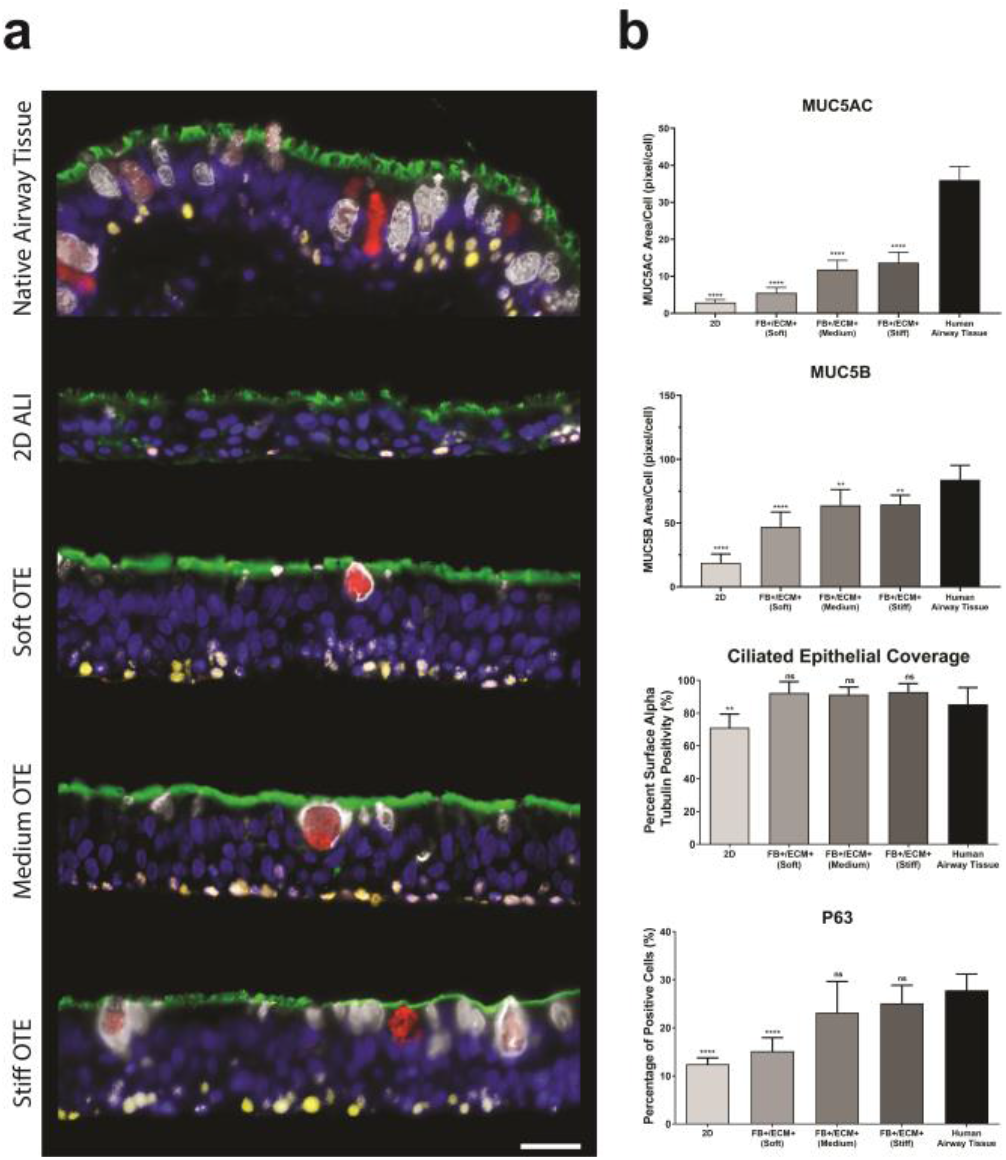
Multispectral Analysis of Epithelial Differentiation Markers. **(A)** Representative multispectral staining of the complete OTE models, 2D ALI culture, and human airway tissue for detection of the following markers: DAPI (blue), Acetylated tubulin (green), MUC5B (white), MUC5AC (red), and TP63 (yellow) (Scale bar = 25μm). **(B)** Associated quantitative analysis of HBE differentiation markers (acetylated tubulin, MUC5AC, MUC5B, and TP63) between the OTE models, standard 2D ALI culture, and human airway tissue. (ns = not significant, ** = p<0.01, *** = p<0.001, **** = p < 0.0001).

### INFLUENCE OF 3D COCULTURE MICROENVIRONMENT ON DIFFERENTIAL EPITHELIAL TRANSCRIPTOMICS

Transcriptomic profiles were developed to evaluate phenotypic differences between the three OTE models, 2D ALI culture, and previously published healthy human airway epithelial tissue^36^. Comparison of the top 10% expressed genes revealed a total of 556 genes shared between all cultures models and the *in vivo* airway epithelium, which included most of the key genes of mature epithelium (**Suppl. Fig. S8, Suppl. Table S1**).

The stiffest OTE model was chosen for a more detailed comparison as it showed to be the most similar of the three OTE conditions to *in vivo* airway epithelium. When comparing the top 10% expressed genes, there were a total of 568 genes in all three datasets, which included key genes of mature bronchial epithelium such as SCGB1A1, SCGB3A1, KLK11, TPPP3, TUBA1A, and SNTN (**Fig. 6A, Suppl. Table S2**). This demonstrates an important overlap in gene expression between *in vivo* airway epithelium, 2D ALI culture, and 3D OTE cultures. This is likely due to the considerable effect and advancement in cell culture media optimization and the ALI maturation methodology^37,38^. Interestingly, we identified 25 genes shared only between the 3D OTE and human airway epithelium, which included several ciliogenesis markers related to mature epithelium such as ZMYND10, RSPH4A, and FANK1^39,40^. However, we also identified 710 genes shared by the 3D OTE model and 2D ALI culture, but not *in vivo* airway epithelium, indicating that both *ex vivo* cell culture methods induce similar genetic phenotypes diverging from *in vivo* tissue.

**Figure 6.**
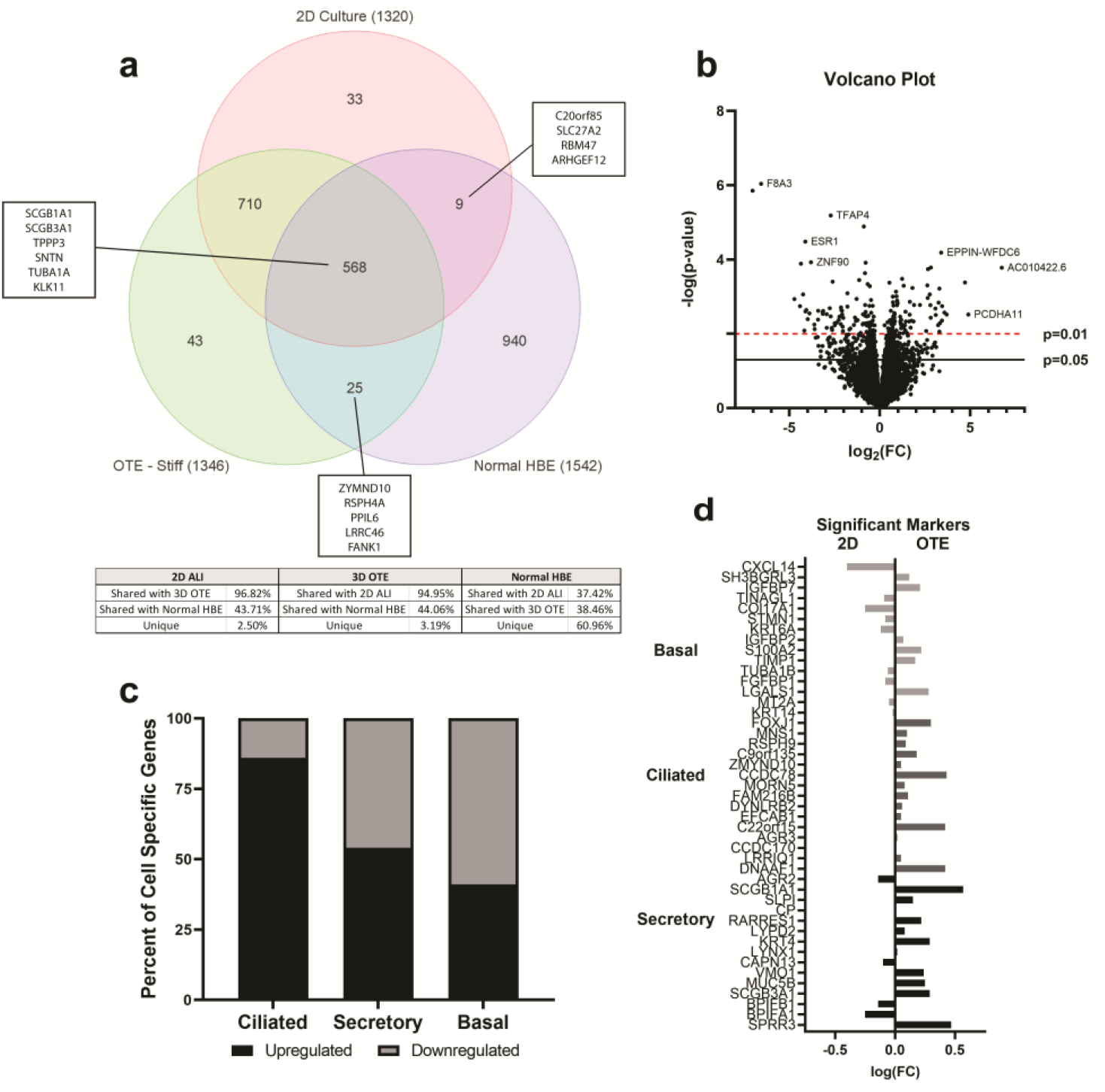
Well-Differentiated HBE Culture Transcriptomics. **(A)** Venn diagram comparing the top 10% gene expression of the stiffest 3D OTE model, 2D ALI culture, and *in vivo* human airway epithelium (Normal HBE)^36^. **(B)** Volcano plot displaying differentially expressed transcripts between the stiff 3D OTE model and 2D ALI culture. **(C)** Expression regulation breakdown of the genes from the gene expression signatures of ciliated, secretory, and basal cells between the stiff 3D OTE model and 2D ALI culture. **(D)** Transcriptomic comparison of the fold change of the top 15 expressed gene markers related to HBE subtypes between the stiff 3D OTE model and 2D ALI culture.

To directly assess the cellular heterogeneity of the HBE transcriptome between the 3D stiff OTE model and 2D ALI culture, we compared their transcriptomic profiles (**Figure 6B**). We identified a total of 719 genes upregulated and 761 genes downregulated in the 3D OTE model compared to the 2D ALI culture (fold change > 1.5). From this dataset, the top expressed markers for the secretory, ciliated, and basal (cycling and non-cycling) cell phenotypes were identified from previous literature for phenotypic grouping^41^ (**Suppl. Table S3**). Goblet and club cell gene expression signatures were combined into a secretory phenotype group due to their similar gene profiles^41^. We found that 85% of the ciliated genes and 54% of the secretory genes were upregulated, while 59% of basal genes were downregulated in the 3D OTE models compared to 2D ALI culture (fold change > 1) (**Fig. 6C**). Expression levels of the top 15 genes of each of the gene sets are illustrated in **Figure 6D**.

Gene pathway differences between the stiff 3D OTE model and 2D ALI model were analyzed using NetworkAnalyst. Significant differences (p < 0.05, fold change > 1.20) between the two groups were evaluated for signaling pathways (Kyoto Encyclopedia of Genes and Genomes [KEGG] and Reactome analysis) along with molecular functions and cellular components (PANTHER analysis)^42^. In total, there were 366 genes significantly upregulated and 333 significantly downregulated in the stiff 3D OTE model compared to the 2D ALI cultures. Using KEGG and Reactome pathway analysis, pathways of interest that were upregulated in the 3D OTE model included those relating to xenobiotic metabolism and tight junctions (**Suppl. Table 4**).

Analysis with the PANTHER database demonstrated upregulation of genes associated with cilia and microtubule binding/motor activity indicating a possible upregulation in cilia function (**Suppl. Table S4**). Meanwhile, there were no specific pathways or components of interest that were downregulated in the 3D OTE model compared to 2D ALI culture.

## Discussion

In this study, we describe the development of a novel 3D OTE human airway model on a biomimetic hydrogel substrate, with incorporated native lung fibroblasts and solubilized lung ECM that can maintain ALI. We evaluated the influence of these biomimetic variables on important phenotypic characteristics that define a well-differentiated human airway epithelium. We found that the OTE model is capable of developing a well-differentiated polarized HBE layer epithelium and provides the versatility to adapt these variables to gain a better understanding of human airway tissue and cellular function *in vitro*.

The inclusion of native lung fibroblasts and solubilized lung ECM were essential components for HBE attachment, viability, pseudostratification and differentiation on the hydrogel substrate. Critical phenotypic and functional characteristics were observed in these complete OTE models: increased pseudostratification, increased TEER, and expression of markers for ciliated, goblet, club and basal cells. Specifically, we report here that the epithelium contains a larger population of p63-positive basal cells, MUC5ac-positive goblet cells, MUC5b-positive club cells, and ciliated cells compared to standard 2D ALI culture after 28 days at ALI. Transcriptomic analysis confirmed that the genetic signature of the 3D OTE HBE layer was characteristic of a mature, well-differentiated epithelium.

One of the main novel components of this OTE is the functionalized hydrogel interstitial layer comprised of thiol-modified HA and gelatin that can incorporate solubilized decellularized human lung ECM. The thiol-modified base structure of our hydrogel provides a covalently linked backbone to minimize undesirable contraction and has shown success in several regenerative medicine applications ^43-45^. We illustrated here that HBE can successfully form a monolayer and mature phenotype after 28 days under ALI conditions on hydrogels fabricated within a stiffness range of 2-10kPA in our 3D multicellular OTE model. Histological analysis of the HBE layer demonstrated that the different stiffness hydrogel substrates affected the level of epithelial thickness. While more work is needed to evaluate this phenomenon, hydrogels matching the stiffness of the proximal or distal airways might be an important driver of HBE phenotype and heterogeneity observed across these tissue locations. Mechanical cues have been linked to the biological patterning and branching morphogenesis of the developing lung^46^. Additionally, while not investigated here, this hydrogel composition can reach stiffnesses as high as 20kPa^31^, approximating the range of fibrotic airway stiffness^47^. Stiffer substrates have shown to promote myofibroblast activation and epithelial dysregulation seen in fibrotic phenotypes. For instance, altered mechanical properties are key characteristics of diseases like asthma and pulmonary fibrosis^48^. Therefore, the successful maturation of HBEs shown here on hydrogels with variable stiffness provide an opportunity to further investigate the influence of the surrounding environment’s biomechanical properties on HBE phenotype.

The second crucial component of our OTE model is the incorporation of tissue-specific ECM to achieve a more physiological relevant biochemical environment. We have previously demonstrated successful isolation of ECM components from our decellularization process and their beneficial effects on the native cell types^31^. Our results indicate that the inclusion of lung sECM was crucial for the formation and maturation of the HBE monolayer on our functionalized hydrogel system. While HBE cells have been cultured on individual ECM substrates, such as collagen^49^ and fibrin^50^, they do not provide the same natural attachment points and biochemical cues that influence biological activity seen in tissue-specific ECM^51^. Moreover, current hydrogel formulations lack the discrete ECM quantities seen in our sECM, such as laminin which promotes epithelial attachment, modulate cell behavior, and stimulate basement membrane generation^52^. Although the mechanisms are not fully understood, our results indicate that airway-specific ECM not only promotes epithelial attachment, but also differentiation on an otherwise unsuitable hydrogel surface. Our future studies will evaluate fabrication of a “defined” biomimetic hydrogel including individually incorporated ECM components at ratios and concentrations identified in our analysis. This may overcome the challenges and resources involved in sourcing and isolating human lung sECM, as well as donor variability. However the evaluation of patient-, tissue location- or disease-specific sECM on OTE phenotype is also an area worth exploring.

A third major component of our 3D OTE model is the inclusion of native lung fibroblasts within the interstitial hydrogel matrix. Similar to the addition of lung sECM, our results highlight the importance of native lung fibroblasts in the formation and maturation of the HBE monolayer on our functionalized hydrogel system. Previous studies have described a symbiotic relationship between the HBE layer and the underlying mesenchyme ^53,54^. This beneficial effect on the HBEs is expected to be mainly based on paracrine signaling since there is minimal direct contact of the fibroblasts with the HBE layer throughout the hydrogel in our model, although we cannot rule this out as a contributor. Our results indicate that the combination of sECM with native lung fibroblasts provided the optimal microenvironment for HBE viability and maturation. This leads us to believe that there is an interaction between the sECM and fibroblasts, and the fibroblasts may reorganize the surrounding ECM into a more favorable environment for HBEs. Within a hydrogel environment, lung fibroblasts have shown to secrete ECM and promote improved attachment of HBEs^32^. Similar to the differential sourcing of lung sECM, we can use pathological sources of fibroblasts to showcase their interdependent relationship with the HBE. The inclusion of native lung fibroblasts in the OTE will be a valuable component for pathological phenotypes where fibroblasts play a major role such as pulmonary fibrosis and COPD^20^.

To evaluate differentiation of the HBE layer, the main functional cell types of the airway epithelium were quantified via multispectral analysis. For each cell type quantified, OTEs demonstrated higher expression when compared to 2D culture, regardless of hydrogel stiffness. For ciliation, MUC5b expression, and TP63 expression in our 3D OTE models demonstrated more similar staining levels to the human airway epithelium compared to 2D ALI cultures. MUC5ac showed a significant increase in expression compared to 2D ALI cultures, but still lower than levels observed in human airway epithelium. A possible reason for the decreased expression of these characteristic markers in 2D ALI cultures is that they have shown to have a greater population of undifferentiated columnar epithelium compared to epithelia recovered from *in vivo* airway brushing^37^. Expression of these markers are important characteristics of well-differentiated bronchial epithelium and are functionally important for normal epithelial function. For example, ciliation expression and mucin production are crucial components of the airways defense mechanism to potentially harmful xenobiotics and gases ^55^. Meanwhile, TP63 expression is a key characteristic of the multipotent capacity of the bronchial epithelial to undergo wound repair ^56^.

We also quantified the transcriptomic changes between the 3D OTE and 2D ALI culture using RNA sequencing. Comparison of the transcriptomic profile of the 3D OTEs with healthy airway epithelial tissue illustrated that epithelium grown on the 3D OTEs is capable of differentiating into a mature epithelium. Not surprising, the 3D OTE cultures shared a large amount of transcriptomic similarity to 2D ALI culture and showed similar genotypic profiles with regards to the different epithelial subtypes. However, supporting findings from the multispectral IHC of functional cell types, 3D OTE transcripts showed slightly increased expression of genes relating to ciliated (FOXJ1+) and club cells (SCGB1A1+/MUC5B+). Interestingly, the gene set correlating to basal cells downregulated in the 3D OTE model compared to 2D ALI culture, in contradiction to the TP63 IHC quantification. This disparity could be explained by different subpopulations of basal and suprabasal cells in the airways and culture samples, or the fact that some club cells have shown to express TP63^41,57^. While TP63 is an identifying marker for progenitor basal cells, research has shown other basal cell subpopulations and suprabasal cells that are negative for TP63, but maintain many other markers, such as KRT16 and MKI67, which were upregulated in our 2D ALI cultures^58^.

The functional pathway analysis identified several regulated pathways of interest between the 3D OTE model and standard 2D ALI culture. Each of the databases provided results that supported greater xeniobiotic metabolism and cytochrome P450 activity in the 3D OTE model, which may indicate a greater population and activity of club cells^57^. This is supported by the upregulation in the SCGB1A1 gene expression and multispectral staining. Previously published analysis also found that 2D ALI culture show decreased expression of pathways related to xenobiotic metabolism when compared to *in vivo* airway tissue^37^. This study postulated that this difference could be explained by the consistent exposure to the external environment by *in vivo* epithelium, but our model indicates that a co-culture 3D ECM microenvironment may also play a role. We also identified upregulation of microtubule activity and cilia expression with PANTHER, supporting our other finds showing a greater degree of ciliation in the 3D OTE culture compared to 2D ALI culture. Interestingly, the pathway and cellular component analysis demonstrated increased tight junction expression in the 3D OTE model. Further investigations in the differences in tight junction formation may provide clarity in the differences in TEER measurements between the 3D OTE model and 2D ALI culture.

In our future studies, we hope to gain an improved understanding of the impact of HBE culture variables we have established here, which includes generation of 3D OTEs using cells direct from tissue isolation to evaluate the expected changes in cell phenotype and gene expression caused by initial tissue culture plastic establishment and expansion. Evaluation of sECM and fibroblast concentrations will be an important next step, as well be further dissecting the exact composition of ECM components most suitable for HBE function. Another interesting next step for analyzing our 3D OTE model will be quantifying cell phenotype gene expression throughout ALI differentiation as gene expression is in constant flux as they differentiate during this culture period ^59,60^. These data will be informative with regards to how the 3D OTE models may alter the differentiation timeline and final HBE heterogeneity.

Our 3D multicellular human airway OTE allows for evaluation of a variety of physiologically relevant parameters that can impact HBE differentiation and function. This provides a novel 3D airway model as a viable alternative to the gold standard of 2D ALI culture, as it allows for more complex and robust experimentation of human airway analogs *in vitro*. While we don’t claim that our model currently represents optimal conditions for HBE differentiation and function, we do propose that the capability to alter and evaluate multiple physiological components, such as cell-cell, ECM-cell and substrate stiffness, will provide a promising platform for further improvements of *in vitro* models of the human airway. The versatility of this model allows for the adjustment of typically inalterable variables in standard 2D monoculture for disease modeling such as: pathological ECM ratios related to diseases like asthma or idiopathic pulmonary fibrosis, fibroblast influence on normal and pathological epithelial behavior, and environmental stiffness influence on epithelial phenotype. This 3D OTE model provides important and physiologically relevant variables for experimentation and opportunities to examine pathological phenotypes and immune responses not capable in current *in vitro* models.

## Materials and Methods

### Lung Extracellular Matrix Fabrication & Analysis

Non-diseased, non-smoking human lung donor tissue was obtained (International Institute for the Advancement of Medicine), decellularized, and solubilized following protocols previously published by our group^31^. The solubilized lung ECM (sECM) was aliquoted and stored at -80°C until further use. The ECM was diluted for colorimetric assays to measure collagen, elastin, sulphated glycosaminoglycan, and hyaluronan content (Biocolor, United Kingdom). Fibronectin and laminin were quantified using the Human Laminin ELISA kit (Abcam, United Kingdom) and Human Fibronectin ELISA Kit (R&D Systems, Minneapolis, MN), respectively. To estimate spectral counts of specific collagen types, the sECM was sent to and processed by RayBiotech (Peachtree Corners, GA) using the Human L3 Glass Slide Array. Additionally, total protein content of the sECM was quantified with a Pierce™ BCA Protein Assay Kit (Thermofisher, Waltham, MA).

### Fabrication of Lung ECM Hydrogel

The thiolated gelatin and hyaluronic acid components from HyStem-HP hydrogel kits (Heprasil and Gelin-S, Lot 18E023, ESI-BIO, Alameda, CA) along with a polyethylene glycol (PEG)-based crosslinker were dissolved in a 1:1 solution of sECM and a photoinitiator solution (P-ECM). The photoinitiator (4-(2-hydroxyethoxy)phenyl-(2-propyl)ketone, Sigma) was dissolved in deionized water to create a 0.1% w/v solution. Briefly, Heprasil and Gelin-S were dissolved in the P-ECM to create solutions of 2% w/v. The three components were mixed in a ratio of 2-parts Heprasil, 2-parts Gelin-S, and 1-part crosslinker. The following crosslinkers were tested for analysis: the linear PEG diacrylate (PEGDA) Extralink crosslinker from the HyStem kit at 1% w/v, and an 8-arm 10kDa PEG Alkyne crosslinker (Creative PEGWorks, Chapel Hill, NC) at 5% and 10% w/v. After encapsulation of the cells, the final hydrogel solution was irradiated with ultraviolet light (365nm, 18w/cm2, BlueWave 75 UV Light Curing Spot Lamp, Dymax, Torrington, CT) for crosslinking.

### Hydrogel Material Analysis

Cell-free hydrogels were prepared, with and without lung ECM, and analyzed for rheology and pore size. The hydrogels were mechanically tested at room temperature with a Discovery HR-2 Rheometer fitted with a steel 8 mm parallel plate geometry. The plate geometry was incrementally lowered into the hydrogel until a normal force of about 0.01N was achieved. To determine the elastic modulus, a compression test was run at a rate of 10 μm/s for a distance of 1.5mm or until the hydrogel fractured, as previously described ^61^. Data were collected every 25ms for stress, force, and gap distance measurements to calculate the strain. The stress-strain measurements were utilized to establish a characteristic curve to calculate the elastic modulus. Additionally, a strain sweep test was run from .02% to 100% shear strain at an oscillation frequency of 1 Hz, during which the storage modulus was recorded, as previously described ^31^.

For pore size, hydrogels were fabricated and allowed to swell in a phosphate buffered solution (PBS). After swelling, the samples were frozen and lyophilized overnight. The lyophilized gels were mounted onto an SEM stub, sputter-coated with 4nm of gold-platinum, and imaged at 5kV using a scanning electron microscope. The SEM images were analyzed with ImageJ (NIH) to quantify pore size.

### Cell culture

Native human lung fibroblasts (Lonza, Basel, Switzerland) were grown in Gibco™ Minimal Essential Media Alpha (Thermofisher, MA, USA) supplemented with 10% fetal bovine serum (FBS), 1% Pen-Strep, and 1% L-glutamine. Primary HBEs (Marsico Lung Institute at the University of North Carolina, Chapel Hill) were cultured in non-proprietary UNC bronchial epithelial growth media (BEGM) on collagen type I-coated dishes. An irradiated fibroblast feeder cell layer and ROCK inhibitor (Y-27632) was supplemented to the culture as previously described ^62,63^. At passage 4, HBEs were utilized for 2D ALI culture and OTE culture. At 70-90% confluency the cells were double trypsinized, counted and seeded on Millicell inserts and OTE cultures.

### 2D ALI Culture

HBEs were seeded on 0.4μm pore size Millicell inserts (Millipore Sigma, PIHP01250, Billerica, MA) coated with collagen-I at a density of 4.15 × 10^5^ cells/cm^2^. The cells were cultured in non-proprietary “UNC ALI” media (Marsico Lung Institute at the University of North Carolina, Chapel Hill), 300μL in the apical compartment and 3mL in the basal compartment. At 100% confluency, the cultures were switched to ALI culture by aspirating the apical media and providing only basal media to the culture. Confluency of the cultures were confirmed with trans-epithelial electrical resistance (TEER) using STX2 chopstick electrodes connected to an EVOM2 voltmeter (World Precision Instruments, FL, USA). The apical surface of the culture was washed with PBS once a week to clear mucus.

### 3D Organ Tissue Equivalent (OTE) Fabrication

The OTE model was fabricated on 8μm pore size Millicell inserts (Millipore Sigma, PI8P01250, Billerica, MA). Briefly, 80μL of the lung ECM hydrogel containing 250,000 lung fibroblasts was pipetted into the apical side of the insert and irradiated with UV light. Two days after UV crosslinking of the hydrogel, HBEs were seeded on the apical side of the hydrogel at a density of 4.15 × 10^5^/cm^2^. The OTEs were maintained in the exact manner as 2D cultures and confluency of culture was confirmed with imaging and TEER. The apical surface of the culture was washed with PBS once a week to clear mucus, if necessary.

### Epithelial Surface Area Coverage

To quantify the attachment and coverage of the HBEs between the groups, the HBE cell layer was imaged in brightfield utilizing an inverted Olympus IX83 microscope. The entire surface of the OTE culture was scanned at 10X and stitched together using CellSens software. The percent epithelial coverage of the surface of the culture was quantified from the stitched images (n=5).

### Trans-Epithelial Electrical Resistance (TEER)

TEER was measured using STX2 chopstick electrodes connected to an EVOM2 voltmeter. For resistance measurements, 300μL of media was temporarily added to the apical side of the inserts. Data is presented as mean +/− standard deviation resistance values (n=12) standardized to insert size (Ω·cm^2^). Statistical analysis was completed to compare the resistance values of the final day of culture.

### Histology and Multispectral Immunofluorescence

After 28 days at ALI, 2D and 3D ALI cultures were washed with PBS, fixed in 4% paraformaldehyde, embedded in paraffin, and 4-μm sections were prepared as previously described ^15^. Additionally, sections from donated lung samples (n=3) were fixed, embedded, and sectioned similarly for staining. Hematoxylin and eosin (H&E) and immunostaining were performed on the sections. MATLAB code was utilized to segment the H&E images to quantify epithelial height.

For immunostaining, the following antibodies were utilized: anti-MUC5AC antibody (1:250; ab212636; Abcam), anti-MUC5B antibody (1:2000; University of North Carolina, Chapel Hill), anti-TP63 antibody (1:300; ab76013; Abcam), anti-acetylated α-tubulin antibody (1:5000; T7451; Sigma). Multispectral IHC staining was performed using the serial staining Opal™ 7-color Manual IHC Kit with the following fluorescent markers: Opal 520, Opal 570, Opal 620, and Opal 690. Images were captured and processed using the Nuance Multispectral Imaging System and CellSens Software on Olympus BX-63 upright microscopes. Individual stains of each marker and their associated fluorophore were imaged and utilized to establish a spectral library. Spectral unmixing of the multiplexed panel was performed by Nuance software, and quantitation of the differentiation markers was calculated for the epithelium. Imaging and quantification excluded the edges of cultures to avoid any edge effect seen from the meniscus in the cultures.

### RNA Isolation, Library Preparation, Sequencing, and Analysis

For RNA extraction and isolation, the surface of the HBE cells was washed with PBS two times prior to RNA collection to remove mucus. The cells were collected, centrifuged, and treated using the Direct-zol RNA Miniprep Plus Kit (Zymo Research, Irvine, CA) to isolate the RNA according to the manufacturer’s instructions. RNA was quantified using the DeNovix RNA Assay Kit (DeNovix^®^, Wilmington, DE), and the RNA integrity was determined with an RNA ScreenTape Kit on the TapeStation 2200 system (Agilent, Santa Clara, CA). cDNA libraries were prepared from RNA extracts with a minimum RNA integrity number (RIN) of 9.5 using the NEXTFLEX^®^ Combo-Seq mRNA/miRNA Kit (PerkinElmer^®^, Waltham, MA). Libraries were quantified using a KAPA Library Quantification Kit (Roche Sequencing and Life Science, Indianapolis, IN) with average fragment length determined by a DNA ScreenTape Kit on the TapeStation 2200 system (Agilent). Finally, cDNA libraries were normalized prior to pooling, and the pooled libraries were sequenced using an Illumina^®^ NovaSeq 6000 system (Illumina^®^, Inc., San Diego, CA), generating an average of 30M 1×101 bp reads per sample. Raw sequence reads were imported into Partek^®^ Flow^®^ analysis software (Partek^®^, St. Louis, MO) for analysis. Cutadapt was used to trim the random barcode (4 bases) from the 5’ end and adapter (AAAAAAAAAA) from the 3’ end of the reads^64^. Bases with a Phred quality score <20 were trimmed, and reads <15 bases in length were discarded. High-quality reads were then aligned and quantified to the hg38 GENCODE reference database using STAR^65^ and an expectation/maximization (E/M) algorithm similar to the one in Xing *et al*^66^, respectively. Noncoding transcripts and protein-coding transcripts with an average read count <10 per sample were removed; the remaining protein-coding transcript-level counts were summed to gene level and normalized with the median-of-ratios method used in DESeq2^67^. Expression profiles of each OTE group and 2D culture were compared to normal HBE profiles using human small airway epithelium transcriptome data previously published^36^. In brief, RNA-Seq data from five healthy nonsmokers was downloaded from the Sequence Read Archive (SRA Accession Number SRP005411) and processed using the same steps detailed above.

A pathway enrichment analysis of the significant protein-expressing transcripts (p<0.05) identified in each OTE group and 2D culture was performed using NetworkAnalyst^42^. Lists were extracted and the resulting gene counts and p-value were included in tables.

### Statistical Analysis

Data illustration and analysis were completed in GraphPad Prism 6 software and illustrated as mean ± SD. For each experiment, an n ≥3 was utilized for each experimental group. Statistical significance was determined with a minimum confidence interval of 95%. Specifically, a one-way ANOVA with Tukey’s multiple comparison post hoc test was used for multiple comparisons, while unpaired t-tests were performed to compare the means for two independent groups. Histological and fluorescent images presented in figures were representative of their corresponding experimental groups.

## Acknowledgements

This work was supported by the National Institute for Health (NIH R01 HL143076), Cystic Fibrosis Foundation (MURPHY17I0) and the Gilead Research Scholars Program (GTS 48599). The authors would like to thank Ryan Szczech, Uma Gandhi, Dipasri Konar, Kathryn Crowell, and other Murphy lab members for their contributions to the early development of the airway OTE design.

## Author Contributions

Conceived and designed the experiments: TL, AA, and SVM. Provided human lung samples for comparison: KO. Performed experiments: TL and UG. Analyzed the data: TL, KDR, KS, SW, and SVM. Prepared manuscript text: TL and SVM. All the authors approved the final draft of manuscript.

## Data Availability

The transcriptomic datasets generated for this study are available in the Gene Expression Omnibus (GSE207931).

## Corresponding Authors

Correspondence to Sean Murphy.

