## Supplemental Figures for "Development of a Novel Air-Liquid Interface Airway Tissue Equivalent Model for In Vitro Respiratory Modeling Studies"

**Supplementary Fig S1: H&E Imaging of Decellularized Lung Tissue.** Comparison of native human lung tissue and decellularized human lung tissue demonstrating loss of cellular components in decellularized tissue, as shown by hematoxylin and eosin (H&E) staining.

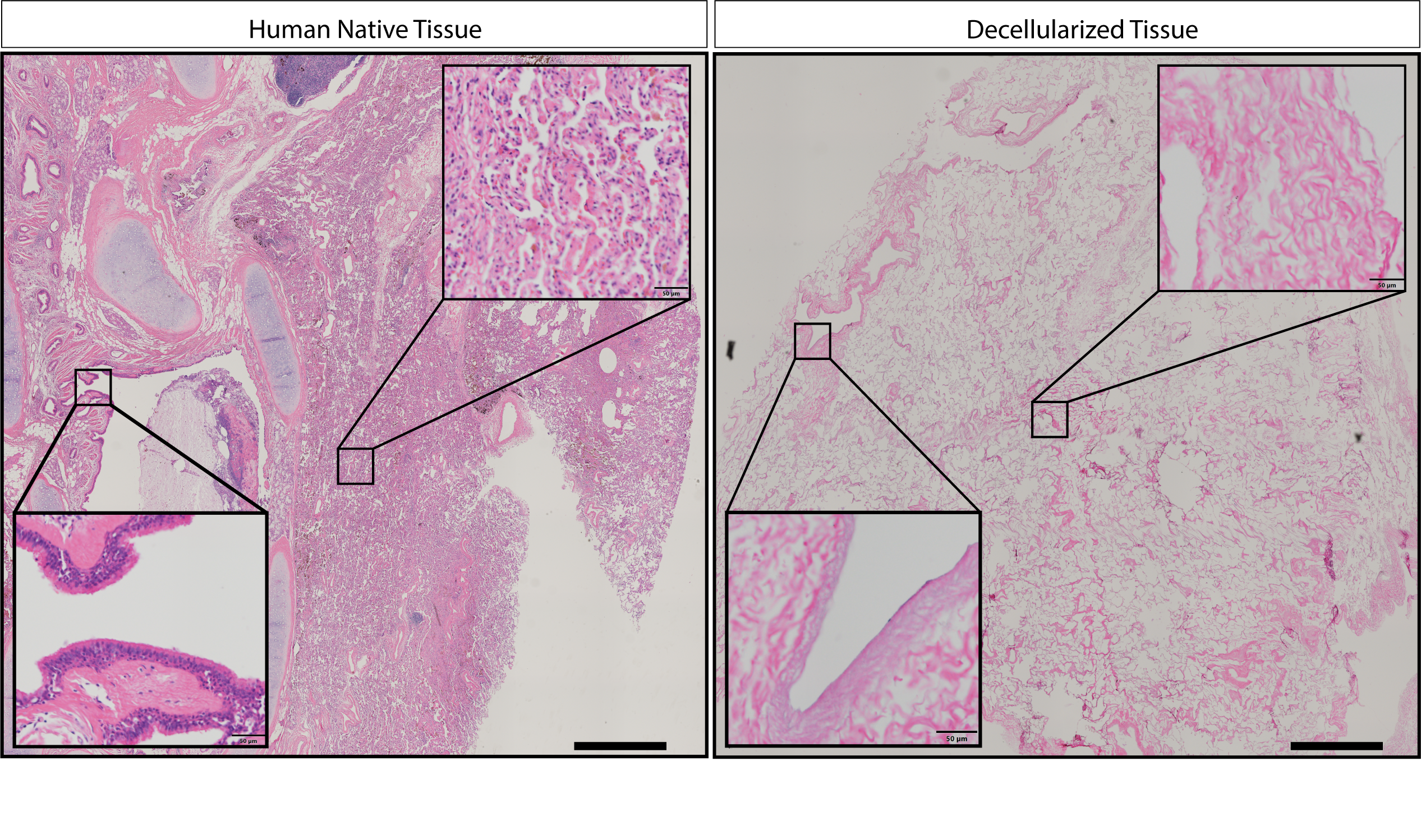

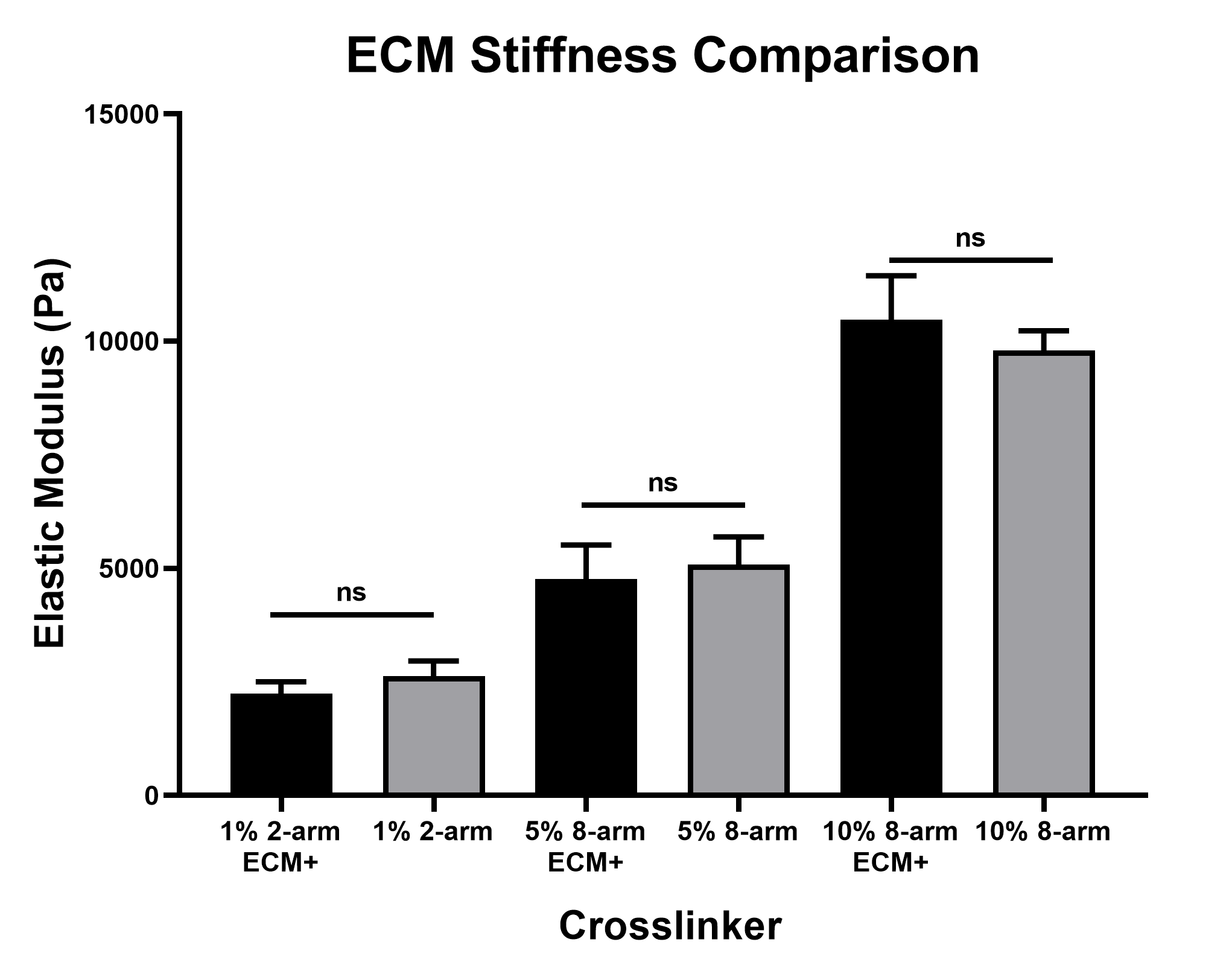

**Supplementary Fig S2: sECM Hydrogel Stiffness Comparison.** Elastic modulus quantification of the three crosslinked hydrogel groups with and without the sECM added into the hydrogel to demonstrate no significant difference in stiffness (ns = not significant).

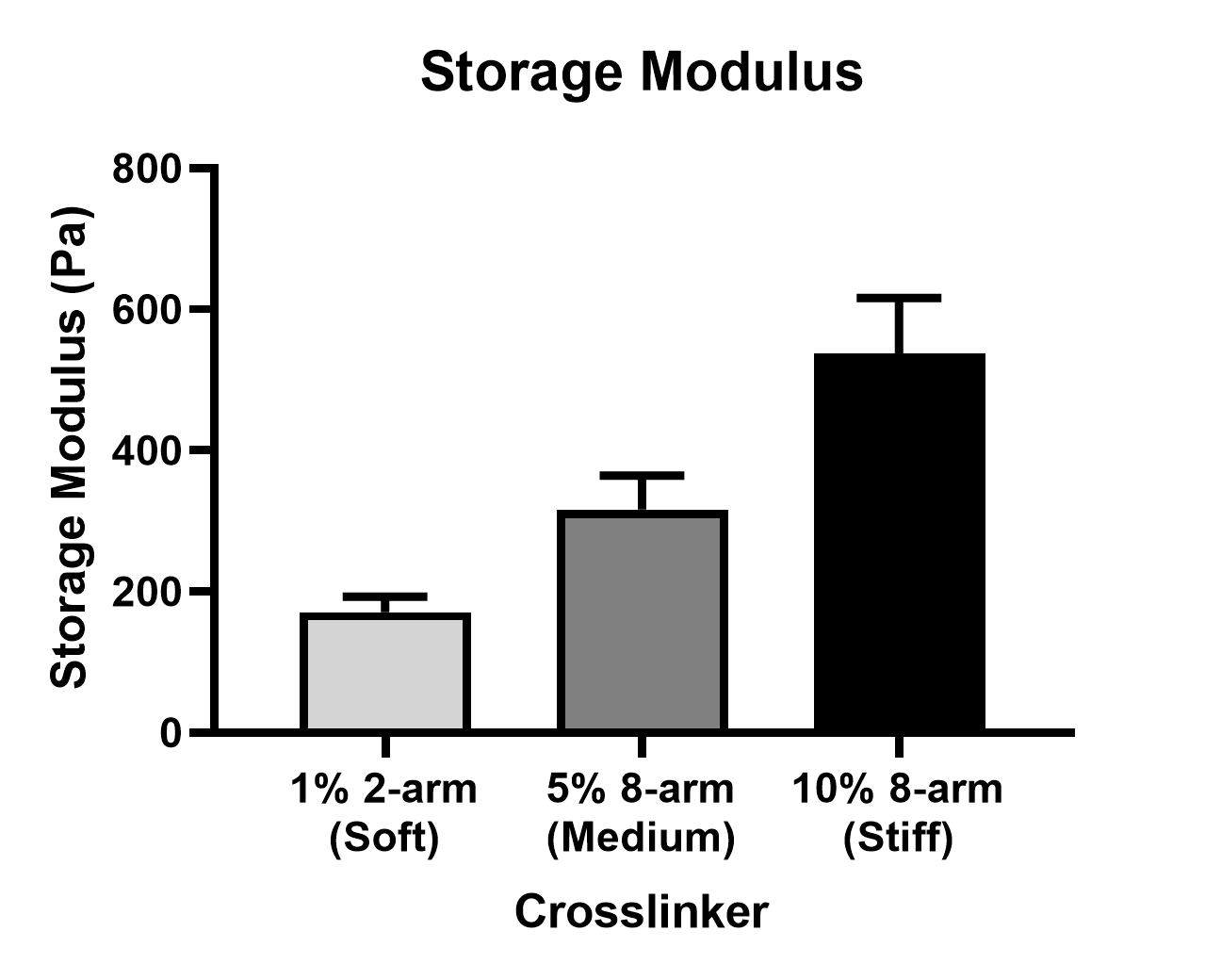

**Supplementary Fig S3: Storage Modulus of Hydrogel.** The storage modulus was quantified for each hydrogel stiffness group and displayed a similar trend of increasing storage modulus across the crosslinking concentrations.

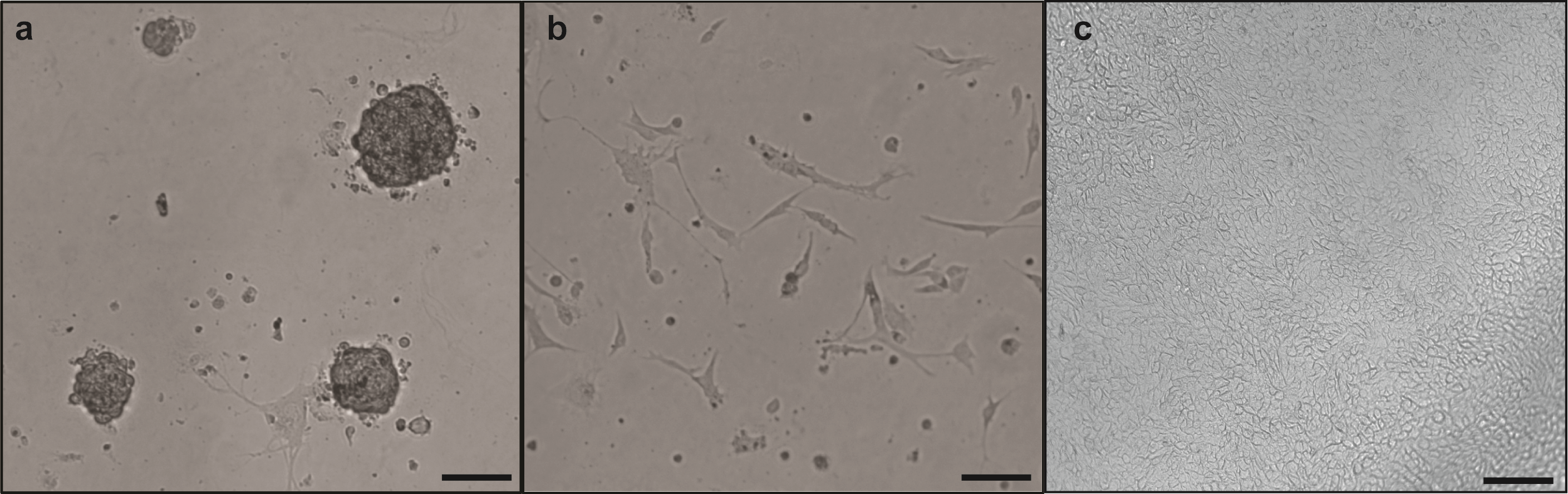

**Supplementary Fig S4: Unhealthy vs Healthy Epithelial Whole Mount Imaging.** Whole mount imaging of the unhealthy epithelial surface of the unsuccessful hydrogel groups showed HBE cells **(A)** attaching to each other in spherical shapes or **(B)** stretching out instead of the typical cobblestone-shaped phenotype. **(C)** Image of healthy confluent monolayer of HBE cells (Scale bar = 100μm).

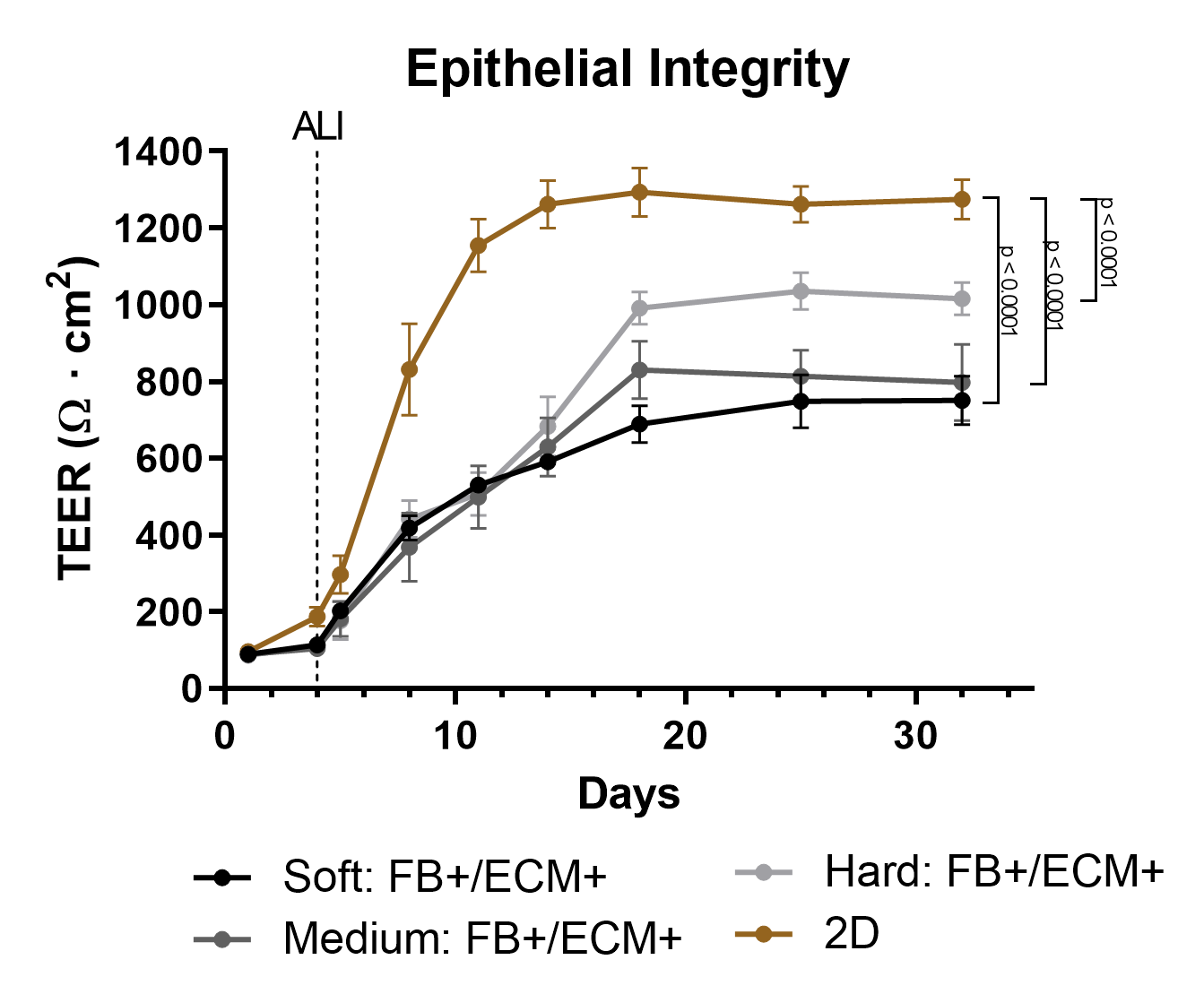

**Supplementary Fig S5: Trans-Epithelial Resistance Comparison.** Trans-epithelial electrical resistance (TEER) was additionally completed on 2D ALI cultures and compared against the 3D OTE models at the three tested stiffnesses. The final TEER value of the 2D ALI group was significantly higher than all three complete 3D OTE models.

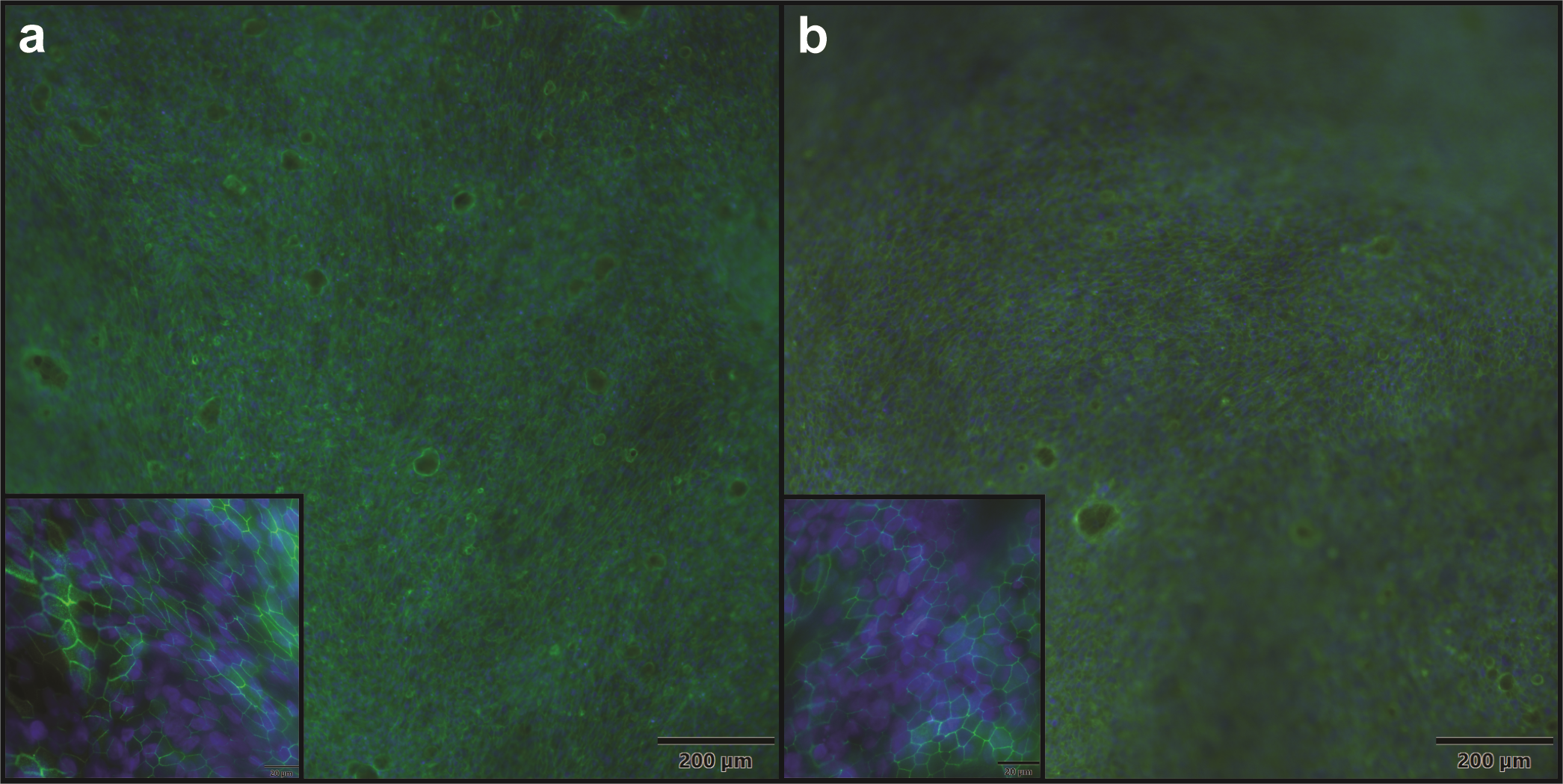

**Supplementary Fig S6: Epithelial Tight Junction Staining.** Whole mount immunostaining of tight junction marker ZO-1 (green) and DAPI (blue) of a representative **(A)** 2D ALI culture and **(B)** complete 3D OTE culture.

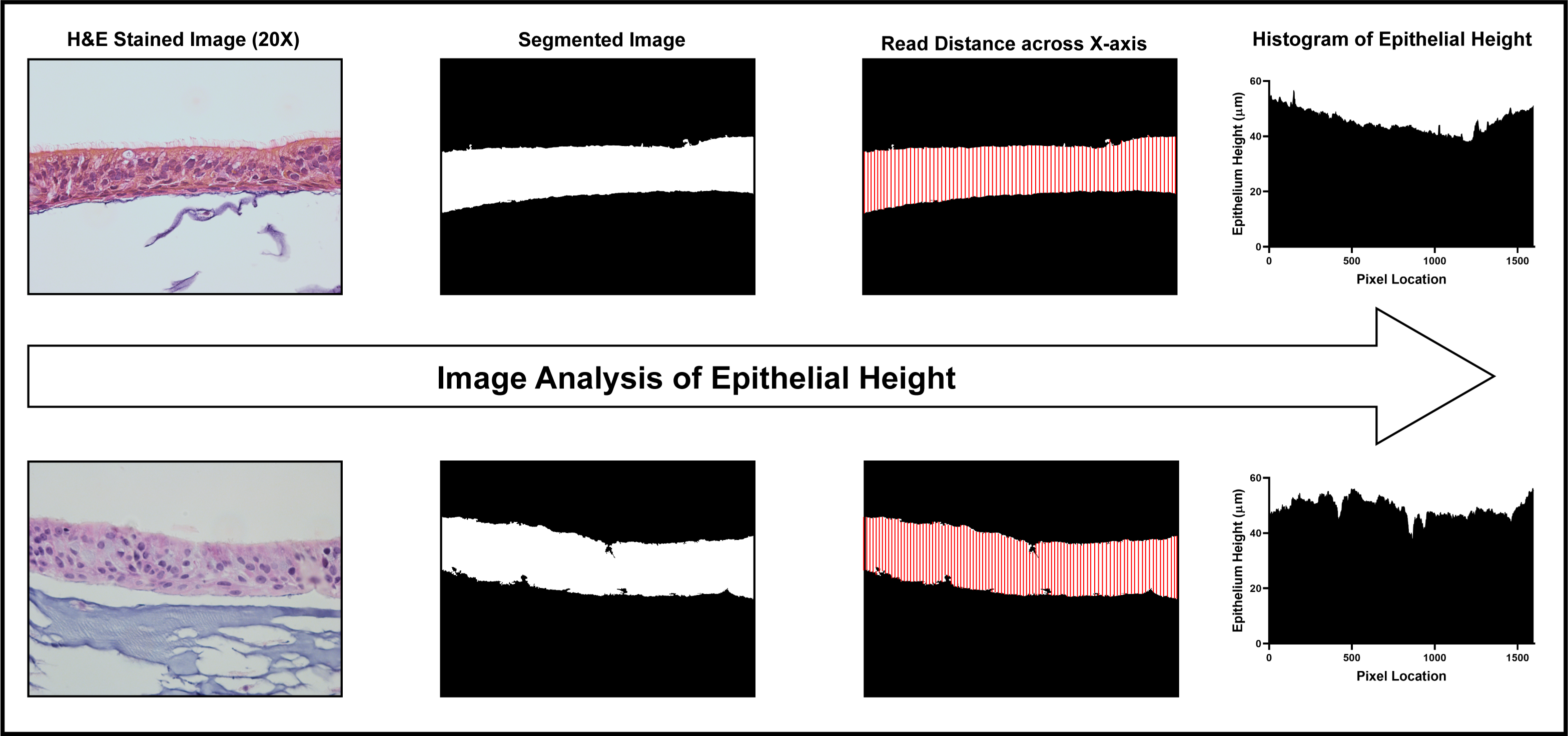

**Supplementary Fig S7: MATLAB Analysis of Epithelial Height.** To quantify the epithelial height of the 2D and 3D OTE cultures along with *in vivo* tissue, H&E images were segmented in MATLAB to isolate the epithelium and the thickness of the area was measured across each x-axis pixel and averaged for a final measurement.

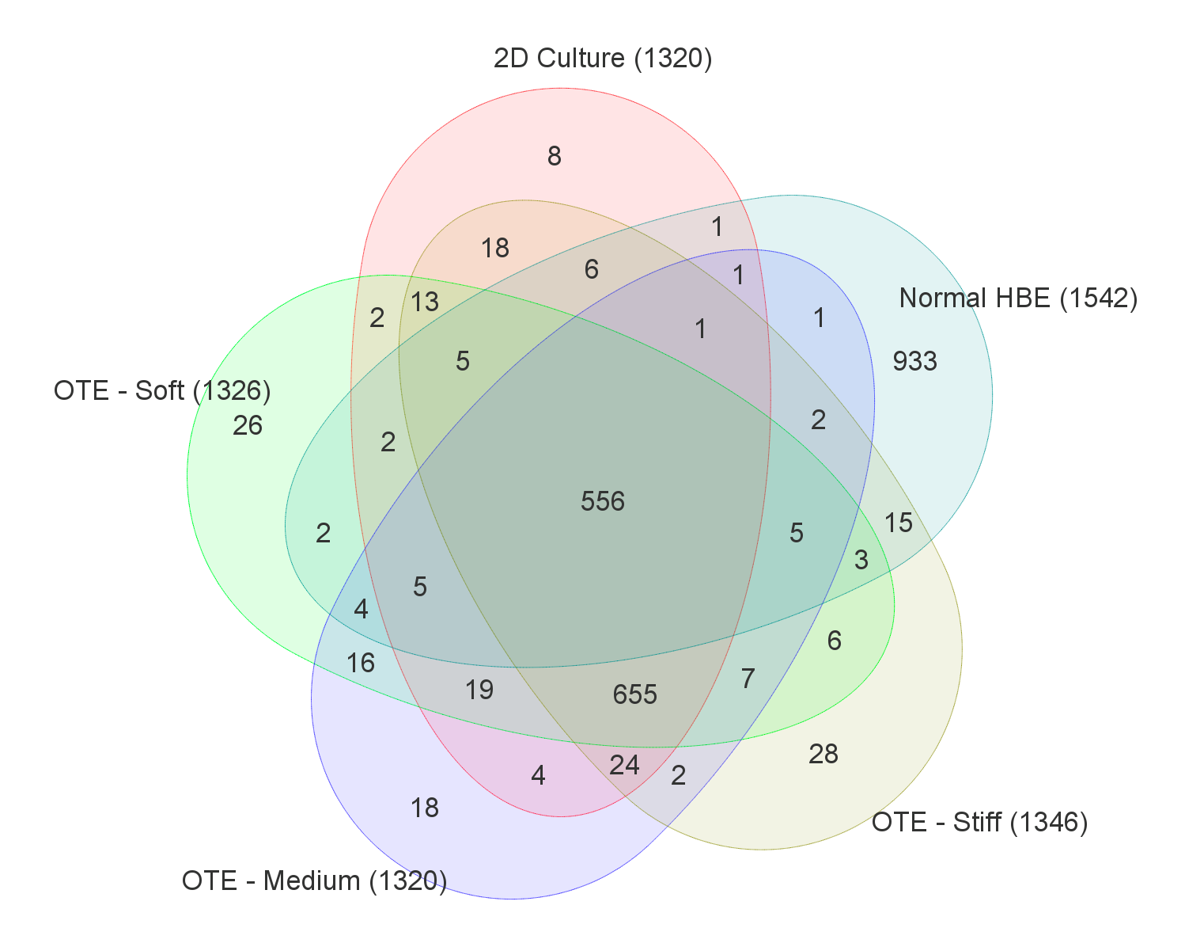

**Supplementary Fig S8.** *Complex Venn Diagram of Top Transcripts.* Venn diagram comparing the top 10% gene expression of the three stiffness 3D OTE models, 2D ALI culture, and *in vivo* human airway epithelium (Normal HBE).
